# Meta-Analyzed Atopic Dermatitis Transcriptome (MAADT) is strongly correlated with disease activity, and consistent with therapeutic effects

**DOI:** 10.1101/2022.05.24.493180

**Authors:** Xingpeng Li, Wen He, Ying Zhang, Karen Page, Craig Hyde, Mateusz Maciejewski

## Abstract

**Background:** Atopic Dermatitis (AD) is a persistent inflammatory disease of the skin to which a few novel treatment options have recently become available. Multiple published datasets, from RNA sequencing (RNA-seq) and microarray experiments performed on lesional (LS) and non-lesional (NL) skin biopsies collected from AD patients, provide a useful resource to better define an AD gene signature and evaluate therapeutic effects.

**Methods:** We evaluated 22 datasets using defined selection criteria and leave-one-out analysis and then carried out a meta-analysis (M-A) to combine 4 RNA-seq datasets and 5 microarray datasets to define a disease gene signature for AD skin tissue. We used this gene signature to evaluate its correlation to disease activity in published AD datasets, as well as the treatment effect of some of the existing and experimental therapies.

**Results:** We report the AD gene signatures developed separately from the RNA-seq or the microarray datasets, as well as a gene signature from datasets combined across these two technologies; all 3 gene signatures showed a strong correlation to the disease activity score (SCORAD) – microarray: Pearson’s *ρ* = 0.651, p-value *<* 0.01, RNA-seq: *ρ* = 0.640, p-value *<* 0.01, combined: *ρ* = 0.649, p-value *<* 0.01. The gene signature improvement (GSI) of two existing effective therapies, Dupilumab and Cyclosporine, as well as that of other experimental treatments, is consistent with their reported cohort level efficacy from the associated clinical trials.

**Conclusions:** The M-A derived AD gene signature provides an evolution of an important resource to correlate gene expression to disease activity and will be helpful for evaluating potential treatment effects for novel therapies.

## Introduction

Atopic dermatitis (AD) is a common chronic inflammatory skin disease characterized by intense pruritus and eczematous lesions [1], with a great unmet need for safer and more effective treatments. AD typically has its onset in infancy or early childhood, showing spontaneous remission in a subset of patients, while others develop a lifelong disease. AD has a significant impact on the quality of life of the patients and represents a significant socio-economic burden [1]. AD is characterized by elevated production of the type 2 cytokines interleukin-4 (IL-4), IL-5, and IL-13, which promote AD pathogenesis [2]. Current treatment strategies in AD have focused on either broad or selective immunosuppression to combat pathologic type 2 inflammation. The approvals of the anti-IL-4R*α* antibody dupilumab, the small-molecule Janus kinase inhibitor baricitinib and abrocitinib, and the anti-IL-13 antibody tralokinumab have provided first-in-class representatives of different therapeutic strategies for the treatment of moderate to severe forms of AD. However, the treatment options remain limited and follow a one-size-fits-all format.

Public transcriptomic data in AD skin biopsies has the potential to serve as a useful resource to better define the AD gene signature and evaluate the therapeutic effects. Meta-analysis has been previously carried out on AD skin biopsy data based on Genechip microarray methods [3]. With the increasing popularity and efficiency of next-generation sequencing (NGS) technology, multiple RNA-seq datasets recorded from AD skin biopsies have been published. Therefore, incorporating these new RNA-seq data into the meta-analysis is desired.

AD transcriptomics data was recorded using microarray technology and published as early as 2006 [4]. A significant contribution to the AD transcriptomics studies was the generation of MADAD (meta-analysis derived atopic dermatitis), an AD gene signature resulting from a meta-analysis of four microarray studies containing 97 samples, with 54 from LS and 43 from NL tissue [3]. MADAD identified a set of 595 DEGs (387 up- and 208 downregulated) using absolute fold-change (|FC|) ≥ 2 and false discovery rate (FDR) ≤ 0.05 as the cutoff criteria. With the development of NGS technology, RNA-seq provided an unbiased assessment of gene expression. It has been reported that RNA-seq outperforms microarray in determining the transcriptomic characteristics of cancer, and performs similarly in clinical endpoints [5]. RNA-seq was also found to be superior in detecting low abundance transcripts in activated T-cells [6]. Benefiting from the improvement in throughput and the decrease of pricing of NGS, RNA-seq has been used in an increasing number of AD transcriptomics studies [7–9]. As a result, it is imperative to develop a new meta-analysis to include both data types and evaluate the AD gene signature from each technology, as well as combined technologies.

Here, we wanted to shed light on the molecular landscape of AD across the publicly available transcriptomics data by comparing lesional and non-lesional AD samples. Although for many diseases numerous -omics datasets are available, AD being no exception, these datasets routinely differ from one another in many key ways, such as the nature of the patients under study (e.g., in one study all patients could be treatment-naïve, in another, they could have previously undergone therapy), whether just one or multiple samples (e.g. a longitudinal collection) are available from each patient in the study, which instrument was used for transcriptomics measurements (the batch effect), and the transcriptomics technology that was employed – microarray, or RNA-seq. All of the above issues, save for the final one (microarray vs RNA-seq data), have been studied extensively, and addressed by either attempting to subtract the variance stemming from inter-dataset differences from the data (most typically using `ComBat` [10], an empirical Bayesian modeling-powered method), or by incorporating them as covariates in the model, most typically in the form of meta-analysis (M-A). Numerous libraries are available for carrying M-A, most noteworthy among them are `metafor` [11] and `meta` [12] for general purpose M-A, and `MetaIntegrator` [13] as an example of a M-A library focusing on -omics data. Here, for our AD analysis, we used `metafor` to extend the M-A approach typically used in transcriptomics by enabling simultaneous modeling of microarray, and RNA-seq datasets; this was achieved by incorporating into the model an additional hierarchical level that describes which of these two technologies a given dataset used. Our model (described by equations 1 through 2) and the associated computational R package called Omics Meta-Analysis (`OMA`) is further described in Materials and Methods.

We used `OMA` to conduct a M-A of 4 RNA-seq studies (GSE121212, GSE137430, GSE141574, GSE157194) and 5 microarray studies (GSE107361, GSE130588, GSE133385, GSE140684, GSE58558), consisting of combined 323 patients. The analysis datasets were extracted from a public repository (Gene Expression Omnibus, https://www.ncbi.nlm.nih.gov/geo), and the pre-processing and analytic procedures were followed in each study as appropriate based on their respective platforms on which they were recorded. A M-A model was used to compare lesional and non-lesional biopsies at baseline, resulting in three lists of DEGs: from microarray, from RNA-seq, and Meta-Analyzed Atopic Dermatitis Transcriptome (MAADT) – a list based on a combined microarray and RNA-seq M-A. We then used the obtained list to evaluate selected investigational and approved therapies to calculate Gene Signature Improvement (GSI) and correlate the GSI to clinical improvement.

## Results

### Dataset selected and coherent genes

We have selected the microarray and RNA-seq datasets based on the selection criteria and leave-one-out (LOO) analysis, as described in Materials and Methods. The final datasets that were retained and used in the M-A have been listed in Table 1.

**Table 1.** Datasets selected for M-A. All 5 microarray studies were generated from the same Affymetrix Human Genome U133 Plus 2.0 Array. The four RNA-seq datasets were generated from Illumina HiSeq instruments, one from 2500, one from 3000, and two from 4000.

| Dataset | Lesional # | Nonlesional # | Reference | Platform |
| --- | --- | --- | --- | --- |
| GSE133385 | 30 | 30 | Pavel et al. 2019 | Microarray |
| GSE58558 | 18 | 17 | Khatti et al. 2014 | Microarray |
| GSE130588 | 50 | 41 | Guttman-Yassky et al. 2018 | Microarray |
| GSE140684 | 31 | 21 | Khatti et al. 2017 | Microarray |
| GSE107361 | 39 | 40 | Brunner et al. 2018 | Microarray |
| GSE121212 | 21 | 27 | Tsoi et al. 2019 | RNA-seq Illumina HiSeq 2500 |
| GSE137430 | 40 | 39 | Ungar et al. 2020 | RNA-seq Illumina HiSeq 4000 |
| GSE157194 | 57 | 54 | Möbus et al. 2020 | RNA-seq Illumina HiSeq 3000 |
| GSE141571 | 39 | 41 | Guttman-Yassky et al. 2020 | RNA-seq Illumina HiSeq 4000 |

### Meta-analyzed transcriptome in Atopic Dermatitis

We used the approach described in Materials and Methods to carry out three meta-analyses: 1) a M-A of 5 microarray datasets, 2) a M-A of 4 RNA-seq datasets, and 3) a simultaneous M-A of 9 datasets, 5 from microarray and 4 from RNA-seq, using a model with an additional hierarchical level used to describe the technology that a given dataset was recorded with (microarray or RNA-seq). The datasets that were used for the meta-analyses are listed in Table 1. The findings from these three meta-analyses are described in detail in Figure 1.C.

**Fig. 1.**
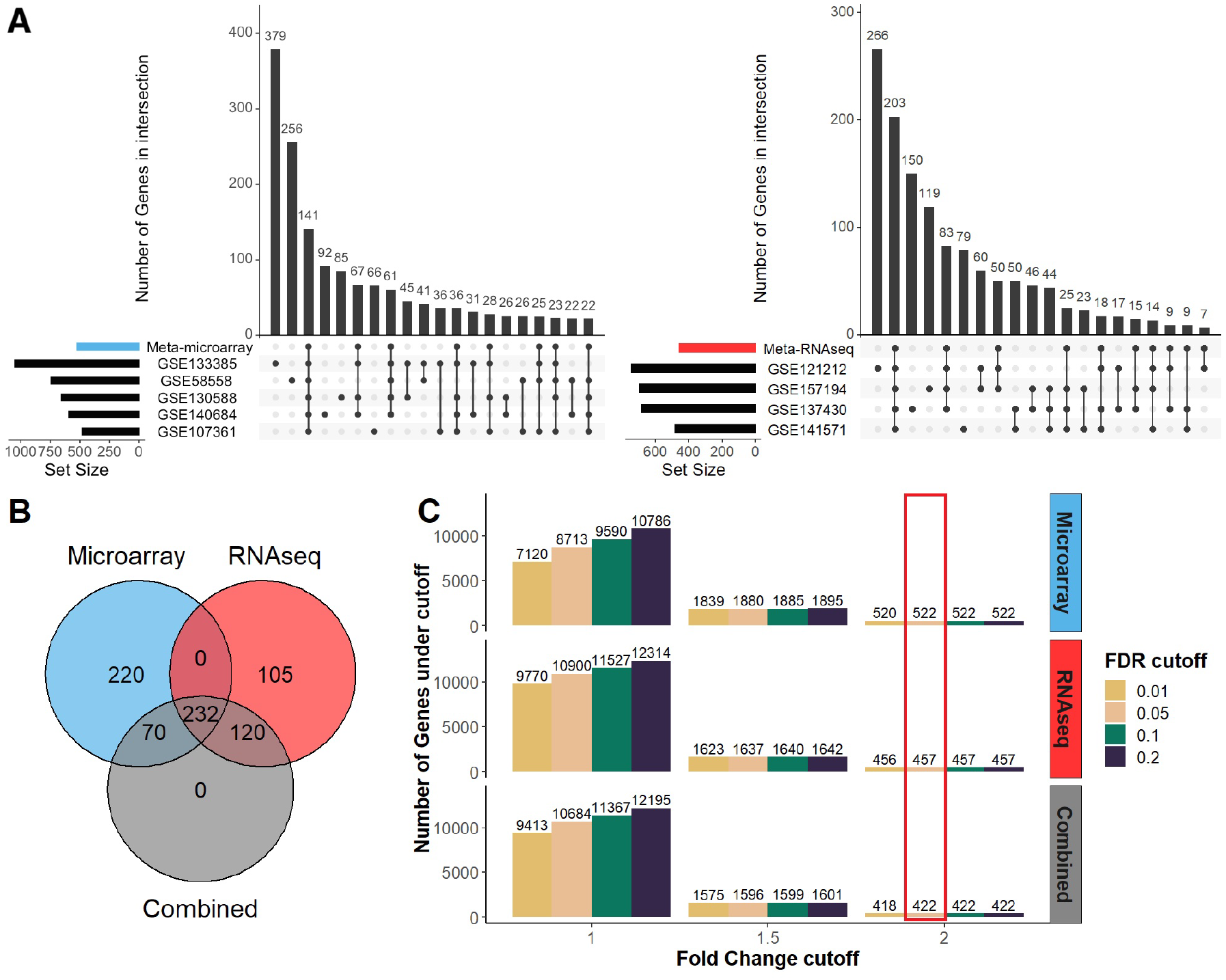
**A** Overlaps of the differentially expressed genes of the individually analyzed datasets and meta-analyzed result under the same threshold (|ES| *≥* 2, FDR *≤* 0.05) with microarray datasets (left) and RNA-seq datasets (right), only top 20 intersect groups were shown for simplicity. **B** Venn diagram of M-A result with RNA-seq, microarray, and combined datasets at same threshold (|ES| *≥* 2, FDR *≤* 0.05) and **C** the number of DE genes from various of thresholds combination for each M-A result, the red box indicates the threshold selected and the number of DE genes for comparison in panels **A** and **B**.

Differential genes can be reported at various effect size (ES) and p-value cutoffs; here, we briefly describe the results from our meta-analyses at the cutoff of |ES| *>* 2, and FDR *<* 0.05. Our microarray M-A brought about 522 differential genes; our RNA-seq M-A resulted in 457 differential genes; the M-A involving both the microarray and RNA-seq datasets resulted in 422 differential genes. A summary of these results can be viewed in Figure 1.C; a full list of results has been provided in Supplementary File 1. Additionally, we have carried out an analogous analysis separately for the significantly underexpressed and overexpressed genes and provided a summary of these results in Supplementary Figure 1.

In Figure 1.B, it can be seen that at our selected thresholds (|ES| ≥ 2, FDR ≤ 0.05), overall, there has been 747 differential genes split between the M-A of microarray datasets, the M-A of the RNA-seq datasets, and the combined microarray and RNA-seq M-A. 220 of these genes (∼29%) were seen in the microarray M-A only. 105 genes (14%) were observed uniquely in the RNA-seq analysis; 190 genes total (25%) were seen in microarray as well as combined M-A (70), or in RNA-seq and combined M-A (120). Finally, 232 the 747 genes (∼31%) were observed in all three analyses.

Among the key AD genes reported in the MADAD study [3], we are able to detect all except IFN. Overall, the results are consistent between MAADT and MADAD. We compared the MAADT list (422 genes) and the MADAD list (594 genes): there are 231 common genes between these two lists, while 191 are unique to MAADT, and 363 are unique to MADAD (Figure 2). We noted that all genes published in MADAD had the same effect size directionality in MAADT (i.e. all significant genes with positive effect sizes in MADAD also had positive effect sizes in MAADT, and those with negative effect sizes in MADAD also had negative effect sizes in MAADT; see Supplementary Figure 2), with an extremely high correlation between the effect sizes in these two studies (Pearson’s correlation of 0.97, p-value *<* 2.2e-16). When we ran the `OMA` analysis on the MADAD datasets, the effect sizes remained highly correlated with MAADT (Pearson’s correlation of 0.82, p-value *<* 2.2e-16), with a high directional concordance (Supplementary Figure 3). Unsurprisingly, key AD genes, including the markers of general inflammation (MMP12), specific T helper activation (e.g. Th2/CCL18, Th1/CXCL10, Th17/PI3/elafin, Th17/Th22 S100A7/A8/A9), and markers of epidermal proliferation (KRT16, Mki67) highlighted in the MADAD paper [3] are also on the MAADT list. Although MAADT and MADAD have a common set of chemokines, which play critical roles in leukocyte migration, there is only one chemokine receptor (CCR7) on the MADAD list, while MAADT has four additional chemokine receptors: CCR1, CCR2, CCR4, and CCR5. To better understand the biology (and biological differences) captured by the lists, we submitted the genes from both lists to GO Biological Process pathway analysis (http://bioinformatics.sdstate.edu/go/) [14]. Results (Supplementary Figure 4) indicate that 231 common genes and 191 genes that are unique to MAADT are mainly enriched in inflammation-related pathways, while 362 genes unique to MADAD are mainly enriched in cell division related pathways.

**Fig. 2.**
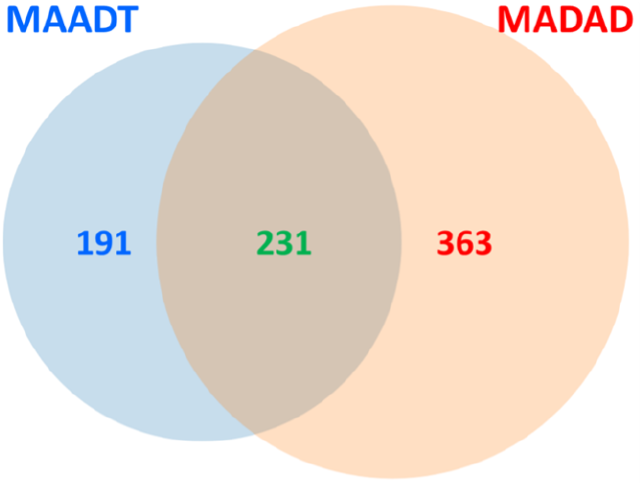
Venn diagram of MAADT and MADAD differentially expressed gene lists. There are 231 common genes between the two lists, while 191 unique to MAADT and 363 unique to MADAD.

**Fig. 3.**
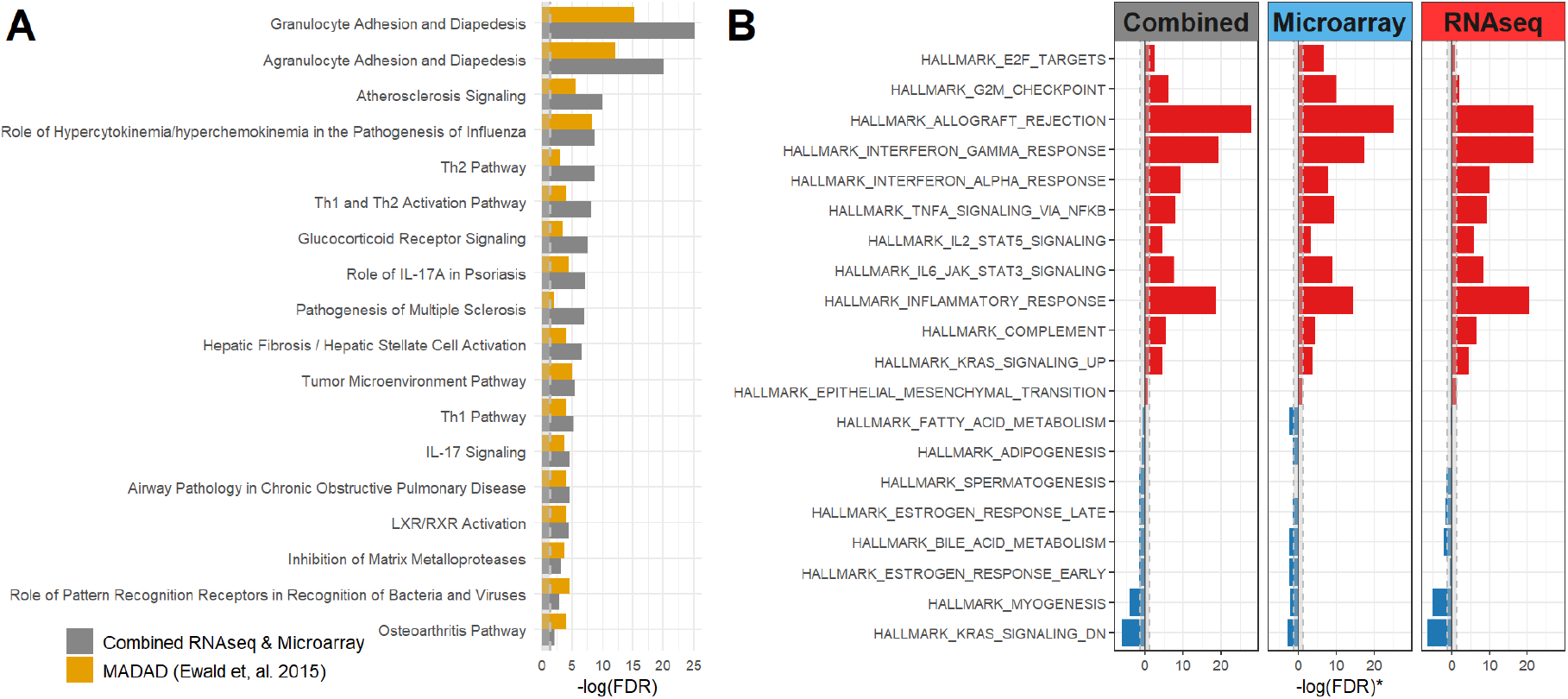
**A** Ingenuity canonical pathways enrichment analysis compared between this M-A combined the RNA-seq & microarray datasets (grey) and the MADAD transcriptome result (yellow) from Ewald et, al. 2015 with the same cutoff at |FC| *≥* 2 and FDR *≤* 0.05. The bars represent the *−*log_10_(FDR), and the grey dotted line shaded area indicates the area where FDR *>* 0.05. The union of top 15 pathways from each gene sets (Combined and MADAD) are shown on the plot. **B** Comparison of pathway enrichment analysis results with hallmark pathways, whereas the bars represent the *−*log_10_(FDR). The sign indicates the enrichment among overexpressed (+) or under-expressed (-) genes, e. g. the positive *−*log_10_(FDR) value indicates the genes upregulated in the pathway are over-represented compared to all upregulated genes in the dataset, and vice versa.

**Fig. 4.**
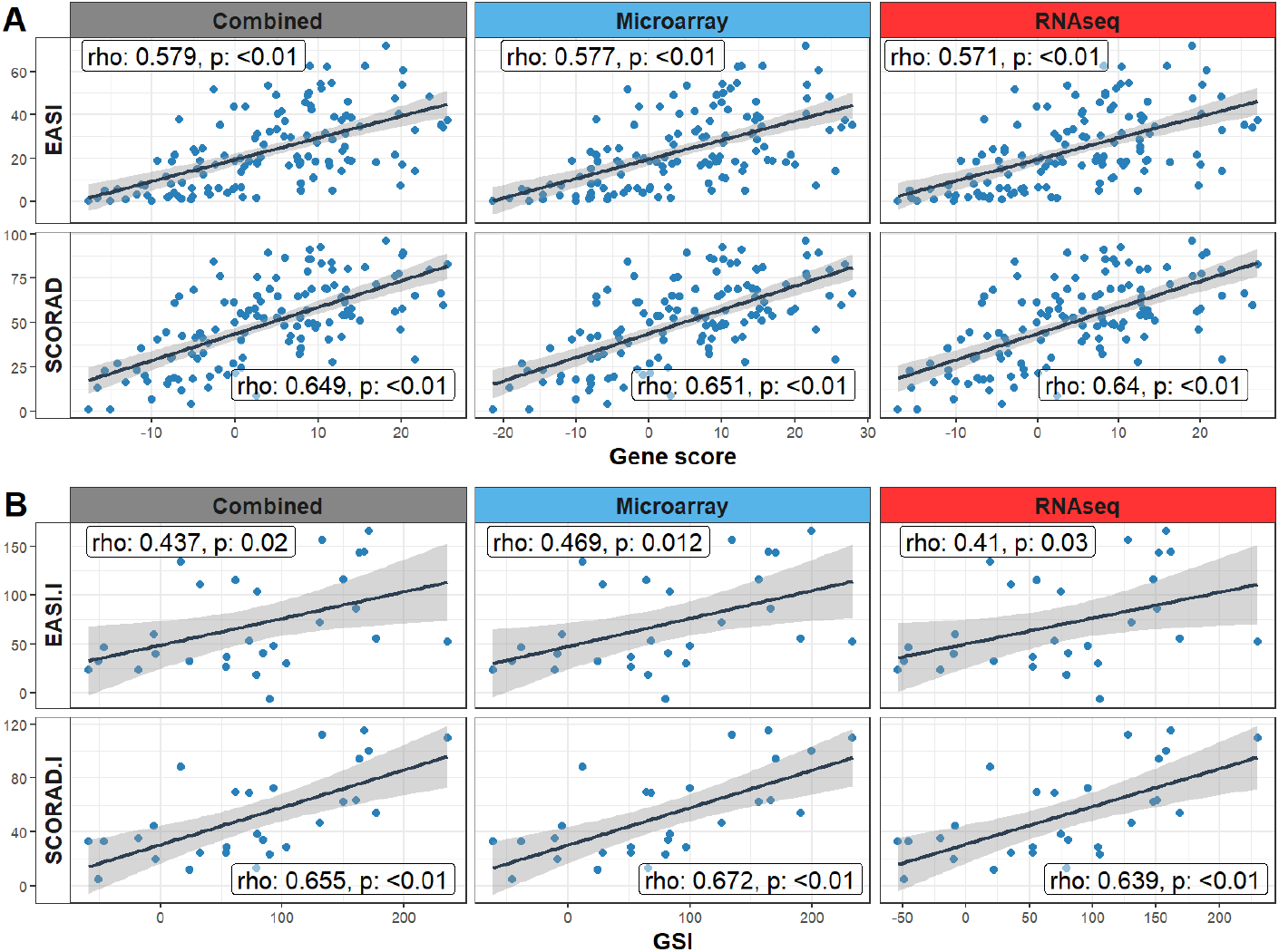
**A** Disease severity index EASI and SCORAD correlation to gene scores from each meta-analyzed result under the same threshold (FC *≥* 2, FDR *≤* 0.05) from published dataset (GSE130588). The gene scores calculated with the z-score method as described in Hanzelmann et, al 2013. **B** Disease severity index EASI and SCORAD improvement relative to baseline (%, y-axis) correlation to Gene Signature Improvement (% GSI, x-axis) with last available time point (Week 16) from published dataset (GSE130588). Only lesional samples at each time point are used for analysis. The correlation coefficient and p-value are calculated with Pearson correlation analysis.

### Ingenuity Pathway Analysis (IPA) and Pathway Enrichment Analysis

IPA was used to identify pathways and functions significantly overrepresented in the transcriptome obtained through M-A. As the pathway information in IPA may have been updated since a previous meta-analyzed MADAD transcriptome from 4 studies by Ewald et al [3], we have re-analyzed the MADAD transcriptome with IPA from the differentially expressed genes (DEGs) reported in MADAD and compared to the result generated from MAAD. The strength of the association of the canonical pathways in terms of −log_10_(FDR) in the MAAD transcriptome was compared to re-analyzed MADAD transcriptome (Figure 3.A).

Among top 15 enriched pathways from MADAD and MAAD transcriptomes, 11 overlap and satisfy the significance threshold of FDR ≤ 0.05. The MAAD transcriptome result yields a more significant over-representation of key immune pathways such as *Granulocyte Adhesion and Diapedesis, Agranulocyte Adhesion and Diapedesis, Atherosclerosis Signaling, Th1, Th2 Pathway* and *IL-17 Signaling*, which are associated with AD [15]. The top two pathways, *Granulocyte Adhesion* and *Diapedesis and Agranulocyte Adhesion and Diapedesis*, both represent the innate immune system where they are involved the process of leukocyte or WBC (White Blood Cell) migration from the blood vessels to the site of pathogenic exposure, which is a key event in the process of inflammation [16]. Atherosclerosis Signaling, which was the third most enriched IPA pathway in MAAD, includes the genes associated with broad vascular inflammation. This is consistent with the previously reported IPA result from the MADAD transcriptome by Ewald et al [3], where the Atherosclerosis Signaling pathway was ranked fourth most enriched pathway. This slight discrepancy could be due to the updated IPA database since 2015.

To further understand the broad biological meaning and the directional changes of the DEGs, 50 hallmark gene sets corresponding to distinct and coherent biological pathways [17] were tested for enrichment from RNA-seq-derived, microarray-derived, and combined DEGs. The positively and negatively regulated genes in each of the three gene sets generated in this study were tested against each set from the collection of 50 above-mentioned sets to evaluate the directional enrichment. There are 20 gene sets that were statistically significantly enriched (FDR ≤ 0.05) in at least one of the three DEG lists generated in this study (Figure 3.B). The enriched upregulated gene sets represented a variety of biological pathways that, again, suggested a close link to immune response such as Interferon Gamma response, Interferon Alpha response, and Inflammatory response. Moreover, the enrichment analysis results from microarray datasets show consistency with the result from RNA-seq datasets. Collectively, the MAAD transcriptome presented here provides a robust AD-specific signal and aligns with the existing known disease pathology.

To compare the biological relevance to the disease between MAADT and MADAD. There are 186 genes identified unique to MAADT and 358 genes unique to MADAD, where have 236 genes that are common between two genesets. Three pathway enrichment analysis shows 7 pathways have statistical significance (FDR ≤ 0.05), which includes inflammatory related pathways like keratinization, signaling by interleukins, and IL10 signaling from 186 genes unique to MAADT. And 25 pathways (FDR ≤ 0.05) from 358 genes unique to MADAD do not contain any clear pathways specific to AD or inflammation. In addition, among 40 pathways (FDR ≤ 0.05) derived from 236 genes that are common between MAADT and MADAD, top pathways are IL4/IL13 signaling, Signaling by interleukins and interferon Alpha/Beta Signaling which are all with high relevance to AD. All 7 pathways from MAADT unique genes are overlapped to the 40 pathways derived from 236 common genes between MAADT and MADAD, where none of the pathways from MADAD unique genes are overlapped to the 40 pathways. This indicates the superior biological relevance of MAADT over MADAD to the disease and general inflammation processes.

### Correlation to disease activity

Three disease gene signatures (DGS) were calculated based on the upregulated genes from our RNA-seq, microarray, and combined meta-analyses. To understand the disease relevance to DGS, we examined the association of disease activity, as measured by EASI activity scores and SCORAD scores, with each DGS. We first evaluated the residual of each DGS, EASI, and SCORAD by adjusting the treatment and time variables, and the normally distributed residuals (Supplementary Figures 5 and 6) warranted the Pearson test to evaluate the correlation coefficient and statistical significance. Baseline SCORAD scores are strongly correlated to all 3 DGS with p-value *<* 0.01 and *ρ* ranging from 0.640 to 0.650 (Figure 4.A), whereas Baseline EASI also correlated with all 3 DGS with p-value *<* 0.01, but with slightly weaker correlation at *ρ* ranging from 0.570 to 0.577.

**Fig. 5.**
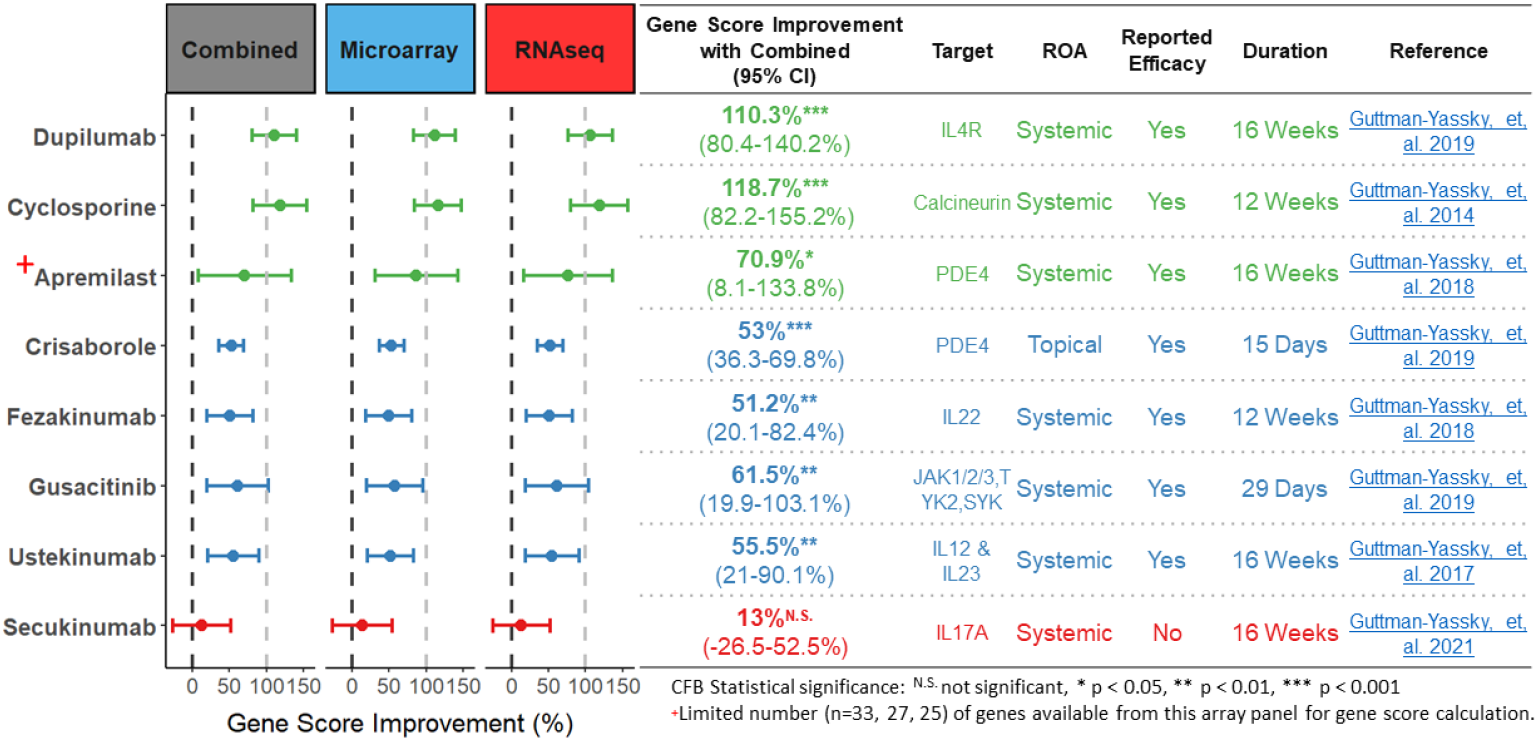
Forest plot with benchmarking result of GSI with RNA-seq, microarray, and Combined AD gene signature on 8 approved or experimental therapies from published datasets. Only one therapy has the route of administration (ROA) through topical treatment. The color on the forest plot indicates the indicates the level of gene signature modulation and the reported efficacy as defined by whether reach statistical significance with primary endpoint at the specified duration.

We also followed individual patients over time to examine whether the change of EASI and SCORAD from baseline was correlated with the gene signature improvement from positively regulated genes with each of the 3 gene sets. Slightly stronger correlations were found between GSI and SCORAD improvement (with all 3 GSI having p-values *<* 0.01 and *ρ* ranging from 0.639 to 0.672), than between GSI and EASI improvement (where the p-values ranged from 0.012 to 0.032 and *ρ* ranged between 0.406 and 0.469, Figure 4.B). In general, DGS derived from either meta-analyzed microarray datasets, RNA-seq datasets, or combined RNA-seq and microarray datasets, all have a stronger correlation to SCORAD than EASI.

### GSI Comparison – investigational and approved therapies

As measured by gene signature improvement through DGS, we compared the GSI with the transcriptome obtained from clinical trials of 7 investigational and 1 approved therapy for AD to evaluate how GSIs (described by equation 3,detailed in Materials and Methods) correlate to the reported clinical efficacy at a specified duration (Figure 5). Dupilumab, which is an IL-4 receptor *α* monoclonal antibody that inhibits the signaling of IL-4 and IL-13, has shown a significant 110.3% gene signature improvement, which is consistent with the reported clinical efficacy in this trial [18]. Cyclosporine displays a 118.7% improvement at the end of week 12, which is also consistent with the positive clinical outcome that was reported in that study [19]. Secukinumab, which is a human IgG1*κ* monoclonal antibody that binds to interleukin-17A, showed no significant gene signature improvement in any of the three gene signatures, and also showed no clinical efficacy in AD from a phase 2 randomized double-blind clinical study [7]. Crisaborole, which is a topical treatment for AD for mild and moderate patients, showed a moderate gene score improvement of 53% with a p- value *<* 0.001. This level of gene score improvement at day 15 is comparable to investigational systemic therapies for AD including Fezakinumab at week 12 [20], Gusacitinib at day 29 [21], and Ustekinumab at week 16 [22].

## Discussion

-Omics profiling, and in particular measuring transcriptional changes between the healthy state and the disease, has become a standard way of investigating the mechanisms pathology. Developments across various -omics technologies over the last decade have been ever accelerating [23], with multi-omics integration increasingly taking the center stage. The ability to integrate multiple datasets has grown in importance and become more commonplace, with numerous M-A methods and studies being published.

AD is a disease that affects numerous individuals across all geographies, therefore understanding its molecular underpinnings has been of high interest and importance. Unsurprisingly, AD was previously investigated via M-A, with MADAD [3] being a key study frequently referred to in pre- clinical inquiries in the pharmaceutical industry. Here, we decided to carry out a new M-A of AD data for several reasons:

- The number of datasets in our study: several datasets have been recorded in the recent years – our M-A consisted of 5 microarray and 4 RNA-seq datasets, while the previous effort consisted of 4 microarray datasets. AD has proven to be a highly heterogeneous disease, and therefore incorporating more datasets in the M-A can help us tease out the signal present across all those suffering from AD, rather than inherent to a specific sub-stratum of the AD population.
- Availability of data from multiple -omics technologies: four of the datasets included in our M-A were recorded using RNA-seq, while previous M-A concerned microarray data only. We were interested in creating a list of genes stemming from the both technologies, thereby mitigating the biases inherent to either microarray or RNA-seq.
- In MADAD, the data was preprocessed using `ComBat` [10], then meta-analyzed using a random effects model. Leaving aside the fact that `ComBat` needs to be applied with care to avoid overcorrecting true biological effects, random effects M-A is meant to account for the batch (or dataset of origin) of a given measurement. Typically, either `ComBat` would be applied to remove the batch effect, or alternatively batch effects would be modeled using fixed or random effect modeling. Removing the batch effects using `ComBat` and then still modeling the batch/dataset of origin as a random effect seems a contradiction.
- Previous methodology favored the random effects model. In our treatment, both the fixed effect and random effect model was considered, and a hybrid result was presented for each gene where a choice was made between the fixed and random effect result based on the amount of excess heterogeneity.
- We wanted to pick the datasets carefully and present here our rejection criteria in a clear fashion. Due to this, although we started with ∼20 microarray datasets, our final analysis utilized only 5 of them, highlighting the importance of scrutiny when using public datasets.

It should be noted that the ability of M-A to increase the focus of the analysis on the signal common to all (or the majority of) the datasets is at the same time a potential limitation of M-A: unless specifically incorporated in the model, subtypes of a disease will not be assumed – the results will focus mostly on the signal that fits a “pan-disease” view of pathology. The fact that we decided to analyze microarray data together with RNA-seq data necessitated developing a slightly altered version of traditional M-A: a hierarchical model with an additional level of hierarchy able to accommodate the influence of the technology with which a dataset was recorded as a contribution to the recorded effect (see Supplementary Table S1).

We have devised three lists of genes expressed differentially between involved AD and non-involved AD tissue: one from the microarray datasets, one from the RNA-seq datasets, and, finally, one from the combined microarray and RNA-seq datasets. As elaborated in the Results section, the overlap between these lists was significant, but there was also a large number of genes differentially expressed only in the microarray list or only in the RNA-seq list. These types of differences should be expected due to the overall number of datasets in each platform still being low, combined to some extent with the systematic biases of each platform. Overall, we recommend using the combined analysis list utilizing the microarray as well as RNA-seq data, due to its sample size being twice as high as either of the other two lists alone, and owing to the fact that, as such, it has the highest potential to reduce both the various technical and sampling biases.

As with all meta-analysis, our study is limited by the sample size of each study included. The 422 MAADT genes missed 220 genes only observed in microarray studies. Some of these 220 genes might be noisier (and missed the |FC| or FDR cutoff) in the RNA-seq datasets; some of these genes might be advantageously selected on the microarray platform, which therefore selectively enriched within the microarray datasets. Similarly, there 105 genes uniquely existed in the RNA-seq platform. The genes presented in MAADT have the advantage of being robustly overexpressed across the majority of the datasets, RNA-seq or microarray, a trend that is desired when combining multiple datasets.

Crucially, the genes present in MAADT (which include genes that are not present in MADAD) correspond to pathways with higher relevance to AD and general inflammation processes, while the genes that are not present in MAADT but are present in MADAD are more in generic biological processes of little significance to AD, which indicates the superior biological relevance to the disease for MAADT.

The disease gene set derived from this meta-analysis showed a strong correlation to the disease activity. A previously reported AD gene set [3] has been widely cited and used to evaluate treatment effects and understand the mechanism of treatment. With additional datasets added to the M-A and the result of gene signature correlation to disease activity score, we presented here an updated AD disease gene set, and provided a resource to evaluate the treatment effects from potential new therapies. In comparison to DE genes derived from single dataset (GSE130588), the gene score or GSI derived from a meta-analyzed gene set has a comparable, if not better, correlation to the disease activity score or disease activity score improvement from that dataset (Supplementary Figure 7). This suggests the derived gene set’s adaptation to other datasets.

In conclusion, we developed a M-A framework for derivation of disease gene sets from multiple datasets, with the ability to handle paired samples, and covariates. We applied this framework to generate an AD gene set for further investigation of AD pathology, biomarker selection, and potential targeted treatment detection. This gene set may serve as an up-to-date reference AD transcriptome for future investigations. The AD genes presented here can be further evaluated through RT-PCR. The transcriptome can also be used to evaluate the response to new treatments, with the help of metrics like the gene signature improvement score used in this study. The framework and the R package developed here can be further used for disease gene set development, of to investigate other pathologies beyond AD.

### Key Points

- We developed a novel method that treats the transcriptomics technology type in each dataset as a separate level of hierarchy in meta-analysis.
- We have used this method to carry out a meta-analysis of 5 microarray and 4 RNA-seq atopic dermatitis datasets and created gene expression and pathway enrichment signatures for this disease.
- Our atopic dermatitis gene signature was significantly correlated with the EASI and SCORAD atopic dermatitis disease activity scores.
- For our gene signatures, we observed a significant gene score improvement in published data demonstrating the effect of multiple approved AD therapeutics – this highlights the practical utility of the gene signature presented here.

## Materials and Methods

### Microarray Dataset Inclusion Criteria and Selection of Coherent Genes

In this study, we have evaluated 16 microarray and 5 RNA-seq datasets (listed in Supplementary Table S1). The list of datasets included for the microarray M-A was built up by sequentially picking a dataset from the list of available datasets with the largest amount of lesional AD samples, checking if including this dataset will increase the overall number of lesional samples in the already selected datasets by more than 5%, then, based on this criterium, either including this dataset or not, and finally discarding the dataset from the list of available datasets; this was repeated until the first dataset that did not meet the “*>*5% criterium” was found, resulting in a list of 7 microarray datasets. Two additional datasets were removed – GSE120899, since it was preprocessed using a batch correction method by the authors and an unprocessed version of the data was not available [24], and GSE99802 was eliminated in the leave-one-out analysis due to its high heterogeneity – resulting in the datasets in Table 1.

### RNA-seq Dataset Processing for Meta-Analysis

The fastq files from four RNA-seq datasets (GSE121212, GSE137430, GSE141571, and GSE157194) were downloaded from GEO using NCBI `SRAToolkit` (https://www.ncbi.nlm.nih.gov/sra), specifically, the compiled binaries/install scripts named “Ubuntu Linux 64 bit architecture”. The SRR numbers or the fastq file names required for the `SRAToolkit` are retrieved from “SRA Run Selector” link in the NCBI GEO accession display page for each GEO number. Processing of the fastq files was performed using the `QuickRNASeq` pipeline [25] utilizing GRCh38 or hg38 for the genome and Gencode v30 for annotation.

### Multilevel Model for Simultaneous Microarray and RNA-seq M-A

The scenario that our modeling approach is concerned with is one in which for a (potentially large) collection of genes *g* we have a number of -omics experiments *s* that span either one or both of the microarray and RNA-seq technologies. Our objective is to, separately for each gene, account for any inter-study and inter-technology heterogeneity, and, for a collection of measured effect sizes derive the true underlying effect size. To do so, we use standard fixed effects and mixed effects methodologies, as detailed below.

M-A of datasets captured by just one of these technologies can be adequately described across all the genes using the fixed effects model:

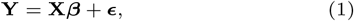

where **X** is the *s*×*f* fixed effects design matrix (*f* is the number of dataset-related fixed effects, and, in our case, we only have a single fixed effect: the expected effect size) common to all genes, ***β*** is the *g* ×*f* matrix of fixed effects, **Y** is the *g* ×*s* matrix of observed effect sizes. ***ϵ*** is the corresponding *g* ×*s* matrix of measurement errors, where for gene *g* and measurement *s* the corresponding error is given by *ϵ*_*gs*_ ∼ *N* (0, *v*_*gs*_), and *v*_*gs*_ is the within-study variance.

This formulation can be extended into a mixed effects model, which additionally captures the random effects:

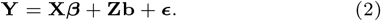

In our current case, we are focused on two specific random effects (inter-study and inter-technology), which are captured by **Z** and **b**:

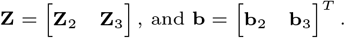

Here, **Z**_2_ is the *s*×*r* _2_ level-two random effect design matrix (where *r*_2_ is the number of level-two random effect classes), **Z**_3_ is the *s*×*r* _3_ level-three random effect design matrix (where *r*_3_ is the number of level-three random effect classes), both common to all genes, and **b**_2_ and **b**_3_ are the corresponding *g* ×*r* _2_ and *g* ×*r* _3_ matrices of level-two (used here to represent the inter-study heterogeneity) and level-three random effects (representing the inter-technology heterogeneity), given by 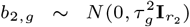 and 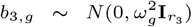, where 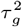 is the inter-measurement variance for gene *g*, and 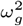 is the inter-technological variance for gene *g*.

It can be seen that if we expand **Zb** in equation 2, we can obtain:

- a standard two-level model (when 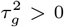, but 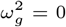 and the **Z**_3_**b**_3_ contribution for gene *g* vanishes),
- a full three-level model with significant inter-dataset and inter-technological variance (when 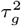 and 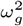 are both non-zero), or
- a model that describes negligible inter-dataset variance in the presence of significant technological variance (when 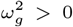, but 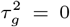 and the **Z**_2_**b**_2_ contribution for gene *g* vanishes).

This three-level approach is also detailed on the `metafor` project’s website^1^, and also in more detail in the core publication referenced therein [26].

In the analyses presented here, we calculate the meta-effect size ***β*** for each gene using both the model represented by equation 1, and the one in equation 2 in the case of two- or three-level M-A (in the case of M-A of effect sizes stemming from two technologies: RNA-seq, and microarray), as described above.

We describe how the model can be fitted efficiently for all genes in the following subsection.

### M-A Model Fitting

In mixed effect M-A, there are several ways to compute the fixed effect estimate 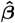 and random effects *τ* ^2^ and *ω*^2^ in equations 1 and 2. Here, we used the Restricted Maximum Likelihood (REML) approach, an efficient method that is known to mitigate the problems that other methods are know to suffer from, for example underestimating the random effects in commonly used methods like DerSimonian-Laird (DL) [27] or Maximum Likelihood Estimation (MLE). In REML, at every step of the optimization loop the fixed effect estimate, 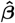, is computed directly, and we maximize the REML ℒ based on that estimate, as well as the current estimates of the random effects *τ* ^2^ and *ω*^2^.

The computation carried out in the REML optimization can be written for every gene *g* as:

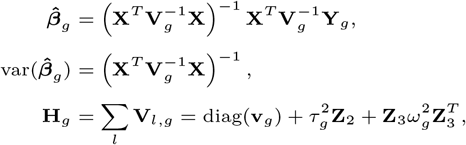

and

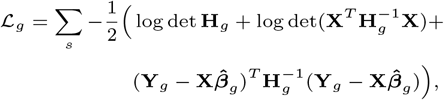

where **H**_*g*_ represents the variance across the three levels *l* considered here, and **v**_*g*_ = [*v*_*g*1_, *v*_*g*2_, …, *v*_*gs*_] is a vector of measurement variances for gene *g*.

### M-A Model Selection

There are various ways to compare and pick between two M-A models, which differ in complexity, that are typically employed depending on what types of models are being compared.

To compare the fixed effect and random effect models, Cochrane’s Q and its corresponding p-value are the standard measures that are typically applied [28] – a practice that we also followed in this work.

To compare two random effect models of varying complexity, the likelihood ratio is a frequently used measure [29]. In our case `L`_*g*_ is measured on the log scale, so when comparing the two- and three-level model, we can calculate the likelihood ratio as:

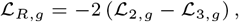

where ℒ_2,*g*_ is the (log) REML for gene *g* obtained from the two-level random effects model (with *τ* ^2^ = 0 and *ω*^2^ ?= 0), ℒ_3,*g*_ is the (log) REML for gene *g* obtained from the three-level random effects model, and ℒ_*R,g*_ is their ratio (expressed as a difference in the log space). For each gene *g*, ℒ_*R,g*_ was compared to the 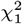 distribution, and we used the resulting p-value to determine whether the more or the less complex model was a more appropriate form of M-A for gene *g*.

### Data Preparation, and Application of DGE and M-A

Each of the datasets listed in Table 1 underwent a final step of preparation before further analysis, where its sample annotation was standardized to Entrez gene IDs. Each dataset was then analyzed separately using `limma` [30], where the lesional samples were contrasted with the non-lesional ones. In the case of RNA-seq datasets, an additional `edgeR` [31] TMM correction was carried out prior to `limma`, followed by normalization (via the `voom` function).

We have carried out three meta-analyses in this study, as described in earlier sections. The two M-A runs, each focusing on a single technology (i.e. the M-A carried out on microarray data only, and the M-A carried out on the RNA-seq data only), were ran using the `rma` function from the `metafor` package. Two meta-analyses were carried out for each technology – one using the fixed effects model, and another using the random effects model (using the REML approach). The p-values of the M-A ES estimate were FDR-corrected [32]. These FDR-corrected p-values along with the ES from the fixed effects and random effects models have been reported here in Supplementary File 1. Additionally, for each gene in the M-A Cochrane’s Q heterogeneity was calculated along with a corresponding Cochrane’s p-value [28]. In each case where the heterogeneity was significant (Cochrane’s Q p-value *<* 0.05), the random effects results were picked; if the heterogeneity was not significant, the fixed effects results were picked; for each gene, these respective picks were presented as the hybrid result (a hybrid of fixed and random effects results). For these hybrid results, FDR-corrected p-values were computed anew using a list of raw p-values from the model selected for each gene.

The mixed M-A where the datasets from microarray were analyzed together with the RNA-seq datasets was carried out using two- or three-level M-A described in detail the previous sections. The results of this analysis were also reported in Supplementary File 1. In this model, we report a two-level result where the technology-based effect *ω*^2^ is reported alongside the ES, and a three-level result where the inter-study heterogeneity *τ* ^2^ was also reported, and a hybrid result that is a mixture of the two former models obtained using the likelihood ratio method described in the previous section. Therefore, the structure of this result in Supplementary File 1 is similar to that of the results from the single-technology analyses, but its interpretation is complicated by the additional level of hierarchy.

### Ingenuity Pathway Analysis and pathway enrichment analysis

IPA software (www.ingenuity.com) was used to characterize pathways significantly over-represented in the identified gene sets following the instruction manual, the DE genes used for IPA analysis was defined with the criteria |Fold-Change| *>* 2 and FDR *<* 0.05. It uses Fisher’s exact test to determine the probability of each biological pathway assigned to each gene set by chance and controlling for the false discovery rate with Benjamini–Hochberg procedure. To identify pathways enriched in the DE genes (|Fold-Change| *>* 1.5 and FDR *<* 0.05) from 3 M-A, we employed the `plot pathway` function in the `RVA` R package for hypergeometric tests and visualization with Hallmark gene sets from the Molecular Signatures Database (version 6.0) [17]. To compare the enriched pathways from the genes unique and common in MADAD/MAADT genes, three pathway enrichment analysis was conducted with `enricher` function from `clusterProfiler` R package using Reactome `geneset` library [33].

### Estimation of Gene Signature Improvement (GSI) scores

Three gene sets from the M-A of RNA-seq, microarray, and combined datasets were quantified by using the z-score method from Gene Set Variation Analysis (GSVA) [34], an unsupervised sample-wise enrichment method that generates a score of activity for gene sets from each sample. Modeling was performed by using mixed effects models, with treatment and time as fixed factors and a random effect for each patient. Fold changes (FCHs) for the change from baseline (CFB) analysis were estimated with the lesional (LS) samples, and hypothesis testing was conducted with contrasts with linear models in R limma package. The averaged baseline gene signature score difference between lesional and non-lesional (NL) skin was calculated. The GSI score 𝒢 was defined as:

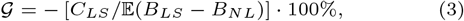

where *C*_*LS*_ is the estimated effect of change from baseline of gene signature score in lesional skin, and 𝔼(*B*_*LS*_ − *B*_*NL*_) is the averaged gene signature score difference of baseline lesional skin *B*_*LS*_ and non-lesional skin *B*_*NL*_. A 𝒢 of 100% indicates a full recovery of lesional skin to the baseline non-lesional level.

### Calculation of GSI correlation to disease activity

The gene expression and meta data along with the disease activity score were obtained from dataset GSE130588 in Gene Expression Omnibus (https://ncbi.nlm.nih.gov/geo). Gene signature scores from positive regulated genes from the 3 gene sets were calculated as described earlier with the z-score method from GSVA. The correlation of disease activity score to gene signature score was evaluated by using Pearson correlation coefficients.

Individual-level GSI was calculated by the difference between post-baseline and baseline gene signature score for each subject over the averaged baseline difference between LS and NL. Two disease activity scores, “Eczema Area and Severity Index” (EASI) and “SCORing Atopic Dermatitis” (SCORAD), were used to calculate clinical improvement. The EASI improvement and SCORAD improvement were calculated as change from baseline over the baseline averaged of each disease activity score. The correlation of disease activity score improvement to GSI was evaluated by Pearson correlation coefficients.

### Statistical Language and Packages

The model used for the multi-level meta-analysis in this study (described in detail earlier in this section) was captured as an R package called `OMA` that we have made available at https://doi.org/10.5281/zenodo.15505102.

## Competing interests

When the work described here was carried out, X.L., W.H., Y.Z., K.P., C.H., and M.M. were full-time employees of Pfizer and owned stock or stock options in Pfizer.

## Acknowledgment

The authors thank the anonymous reviewers for their valuable suggestions. This work is supported in part by funds from the National Science Foundation (NSF: # 1636933 and # 1920920).

## Supplementary Materials

Supplementary materials available online:

- Supp. File 1. Full MAADT result table.
- Supp. Table S2. Meta-analyzed AD genes from LS vs NL with combined RNA-seq and microarray datasets.
- Supp. Table S3. Meta-analyzed AD genes from LS vs NL with RNA-seq datasets.
- Supp. Table S4. Meta-analyzed AD genes from LS vs NL with microarray datasets.
- Supp. Table S5. Ingenuity canonical pathway analysis result of meta-analyzed AD genes from LS vs NL with combined RNA-seq and microarray datasets at cutoff threshold (|FCH| ≥ 2, FDR ≤ 0.05).
- Supp. Table E6. Ingenuity canonical pathway analysis result of meta-analyzed AD genes from LS vs NL with RNA-seq datasets at cutoff threshold (|FCH| ≥ 2, FDR ≤ 0.05).
- Supp. Table E7. Ingenuity canonical pathway analysis result of meta-analyzed AD genes from LS vs NL with microarray datasets at cutoff threshold (|FCH| ≥ 2, FDR ≤ 0.05).

## List of acronyms or abbreviations

## NOMENCLATURE

MAADT: Meta-Analyzed Atopic Dermatitis Transcriptome
AD: Atopic Dermatitis
LS: Lesional
NL: Non-Lesional
M-A: Meta-Analysis
OMA: Omics Meta-Analysis

---

**Supp. Fig S1.**
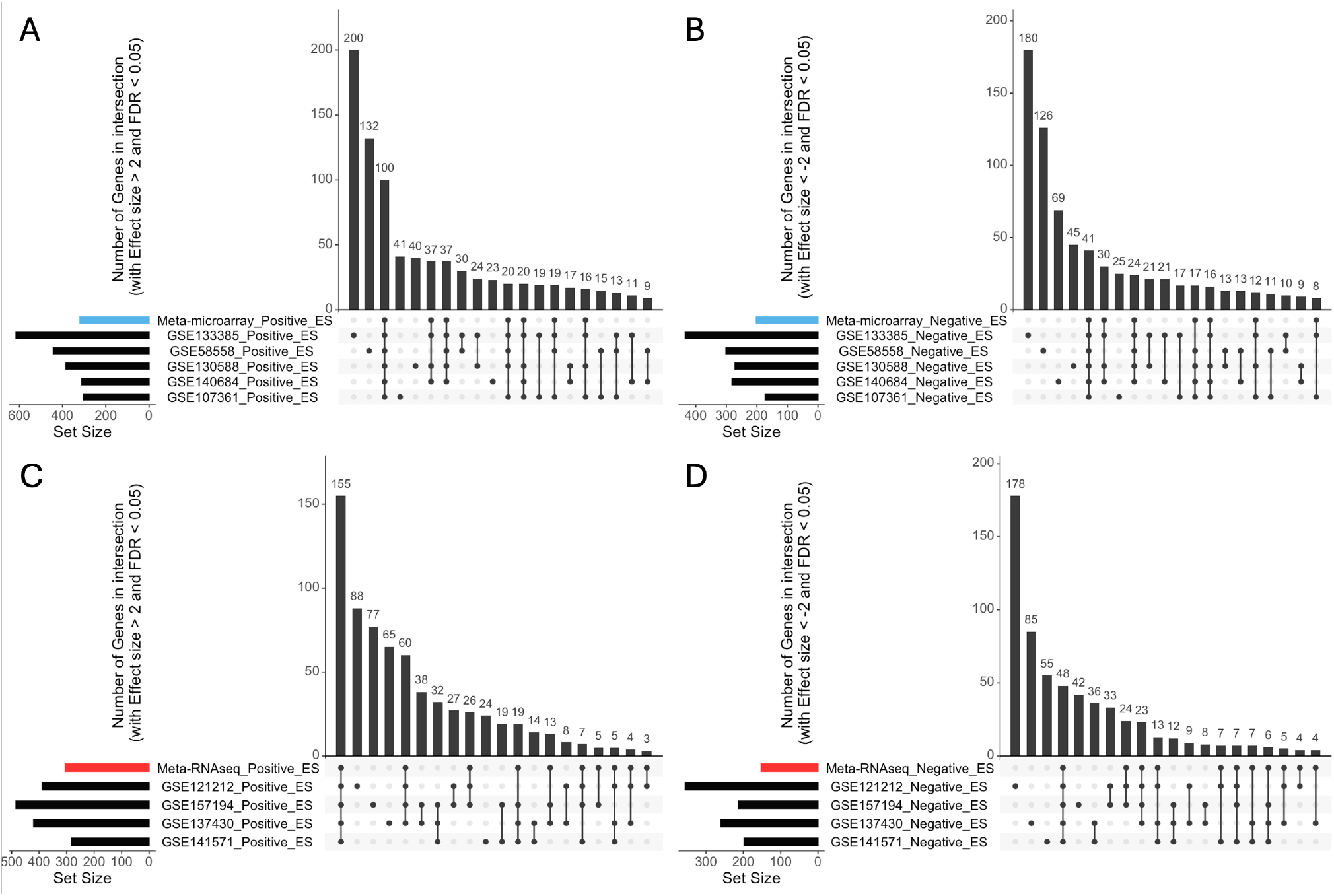
Overlaps of the differentially expressed genes of the individually analyzed datasets and meta-analyzed result under the same threshold with microarray datasets (**A**. ES *≥* 2, FDR *≤* 0.05; **B**. ES *≤* -2, FDR *≤* 0.05) and RNA-seq datasets (**C**. ES *≥* 2, FDR *≤* 0.05; **D**. ES *≤* -2, FDR *≤* 0.05), only top 20 intersect groups were shown for simplicity.

**Supp. Fig S2.**
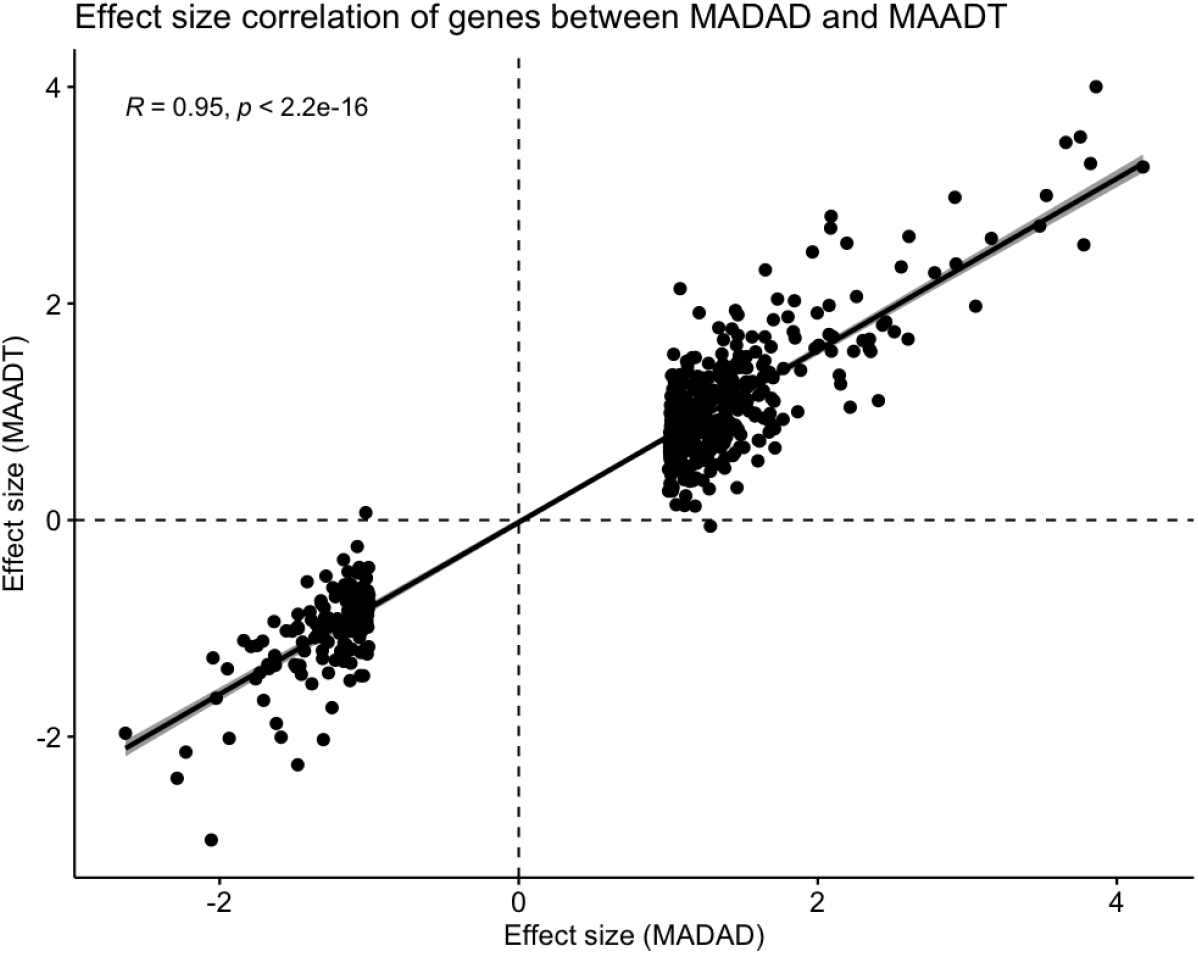
Strong correlation of meta-analyzed effect size between MADAD and MAADT (|FC| *≥* 2, FDR *≤* 0.05).

**Supp. Fig S3.**
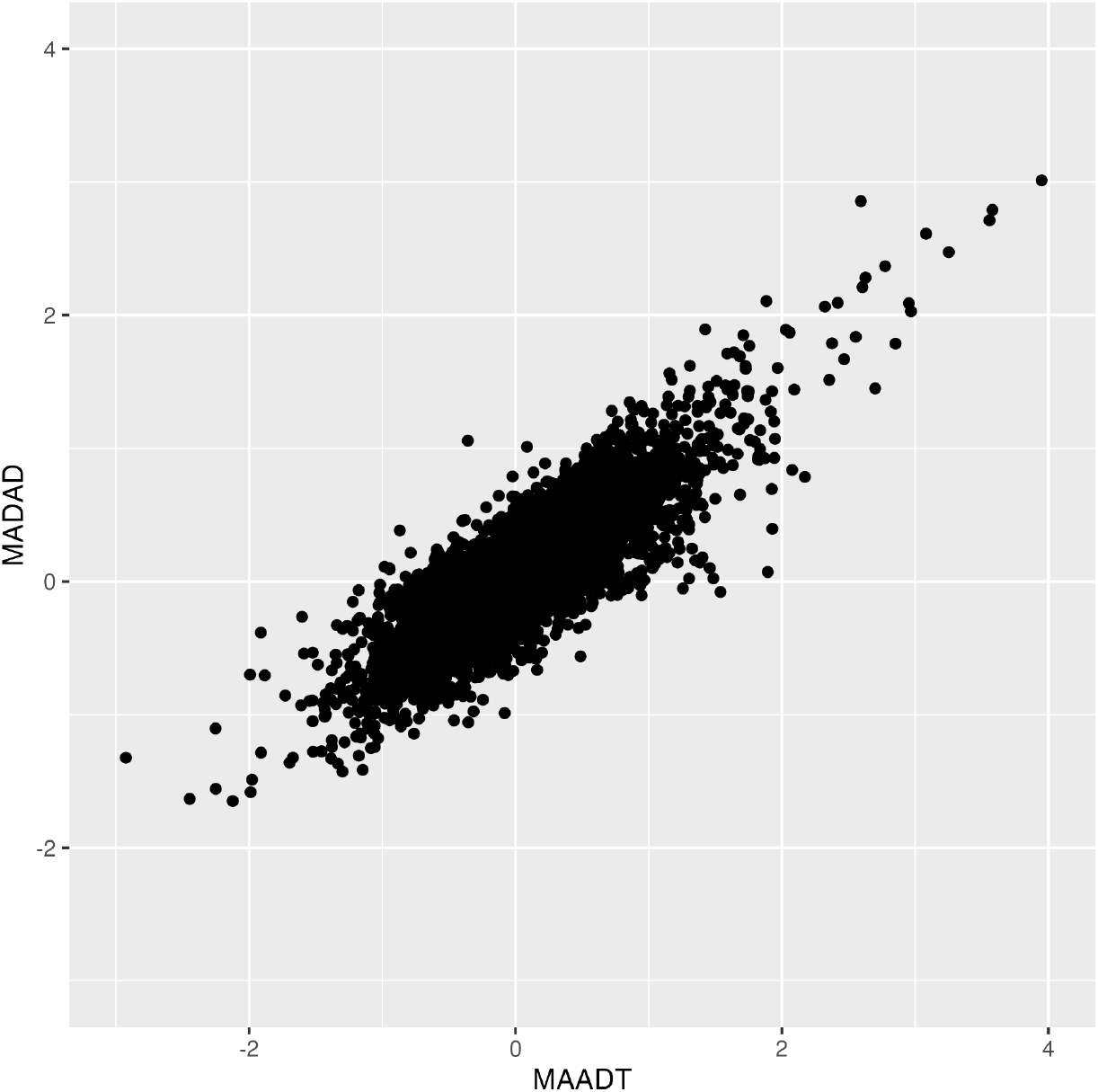
Strong correlation of meta-analyzed effect size between MAADT and MADAD datasets reanalyzed using OMA. Parson’s correlation *ρ* = 0.82, p-value *<* 2e-16.

**Supp. Fig S4.**
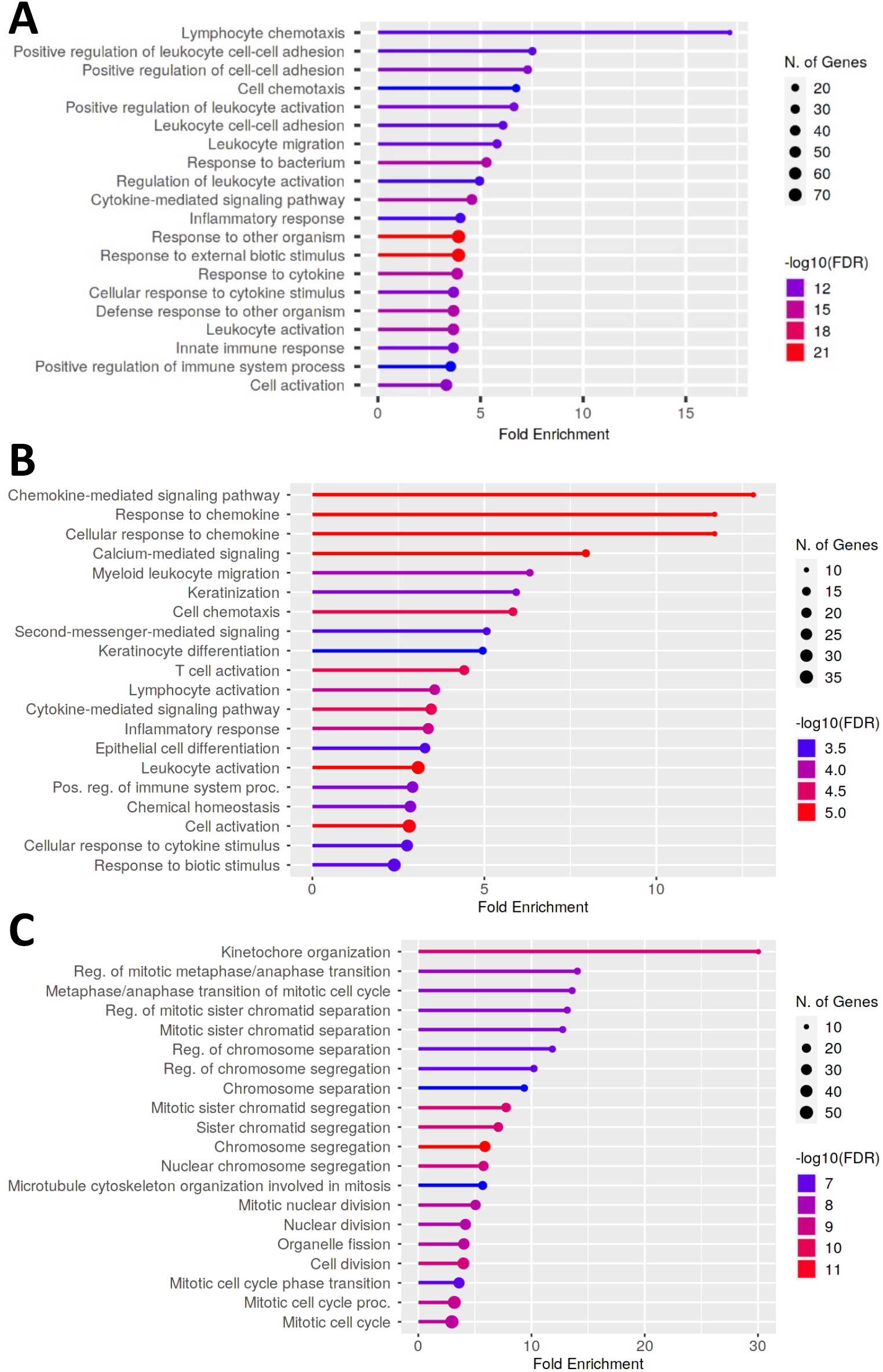
A. Top 20 Go Biological Process pathway among 231 genes common to MAADT and MADAD. **B**. Top 20 GO Biological Process pathway among 191 genes unique to MAADT. **C**. Top 20 GO Biological Process pathway among 362 genes unique to MADAD.

**Supp. Fig S6.**
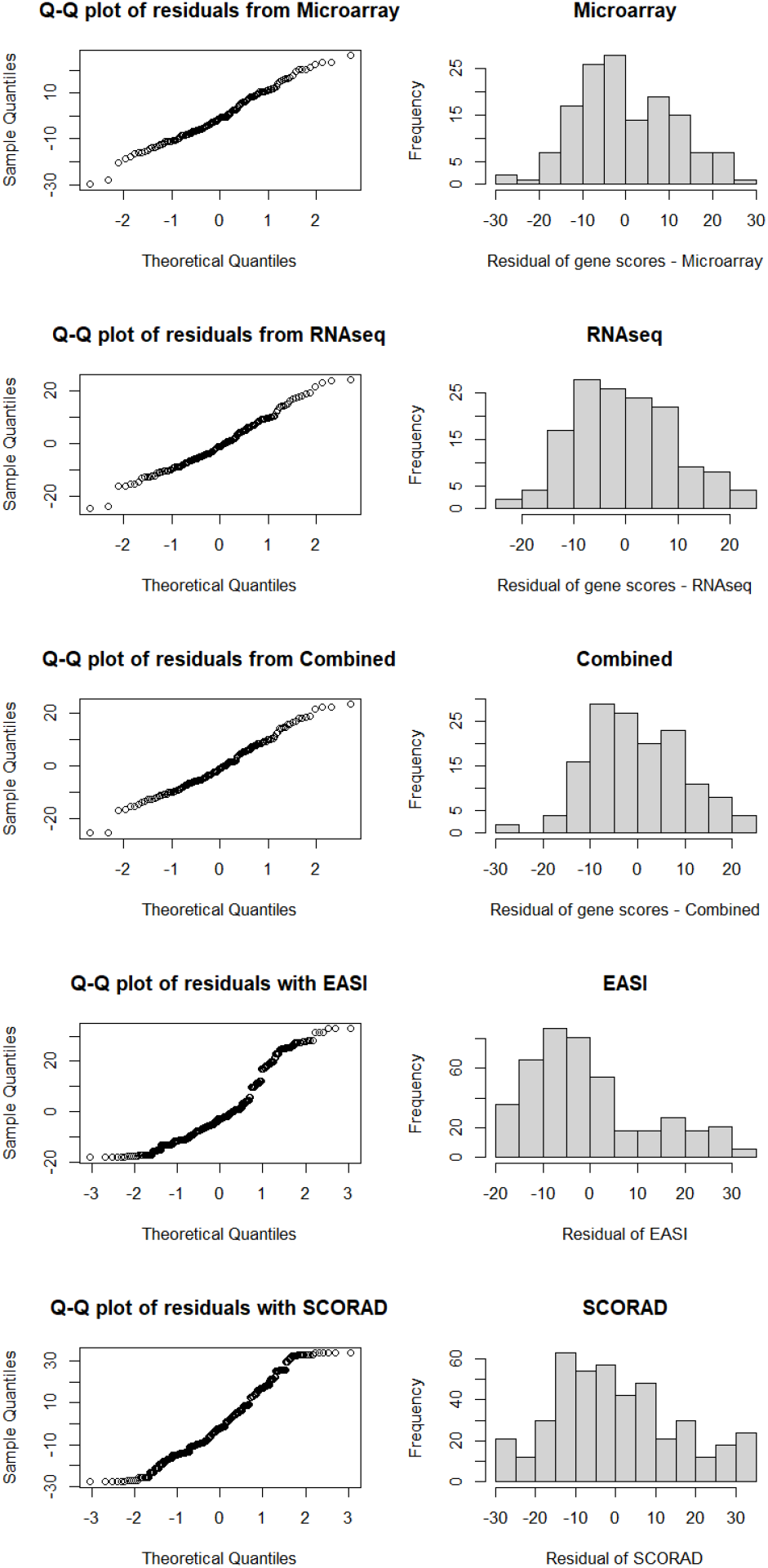
QQ plot and histogram of residuals for EASI, SCORAD, and each gene score *improvement* in dataset GSE130588.

**Supp. Fig S5.**
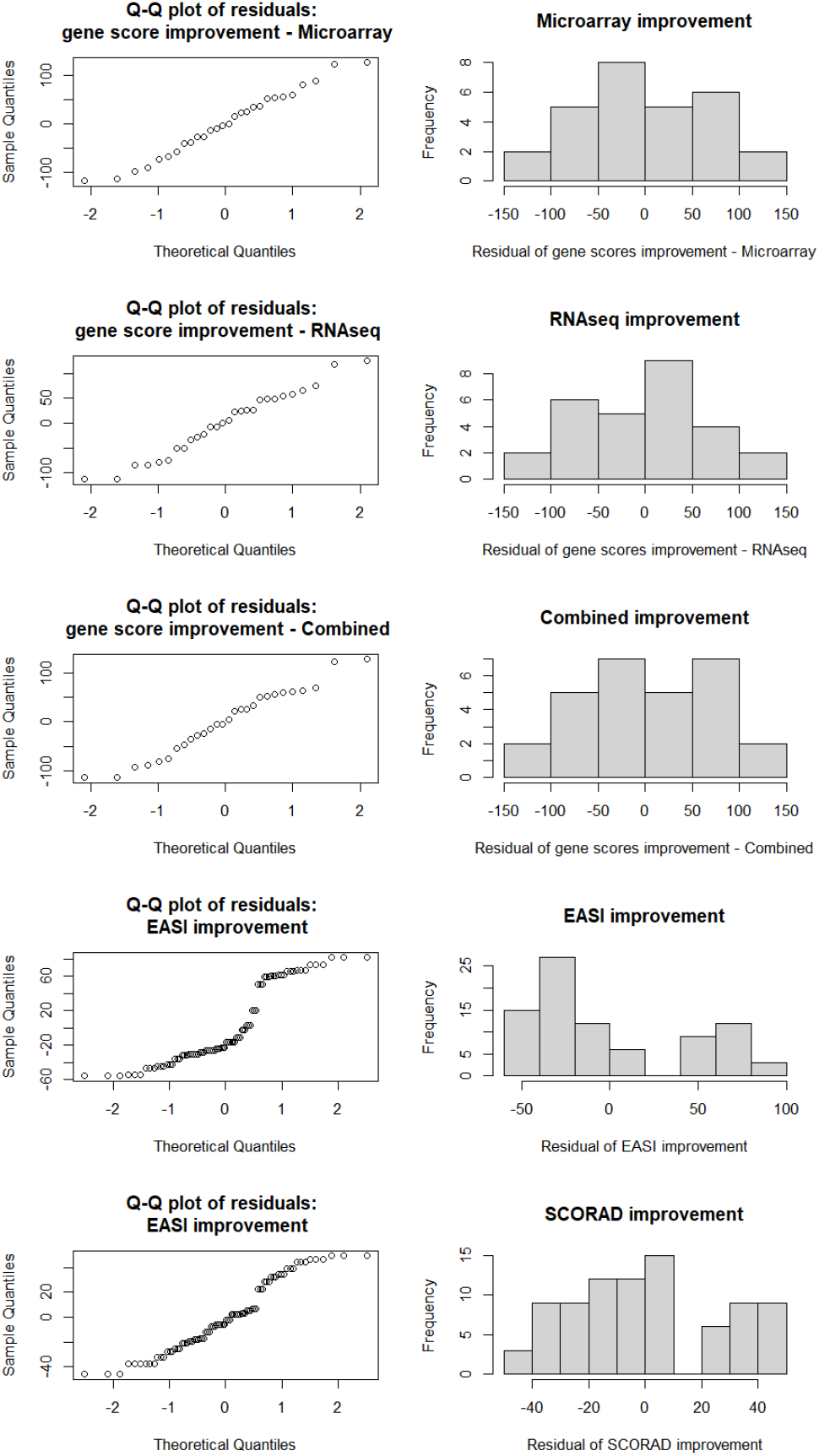
QQ plot and histogram of residuals for EASI, SCORAD, and each gene score in dataset GSE130588.

**Supp. Fig S7.**
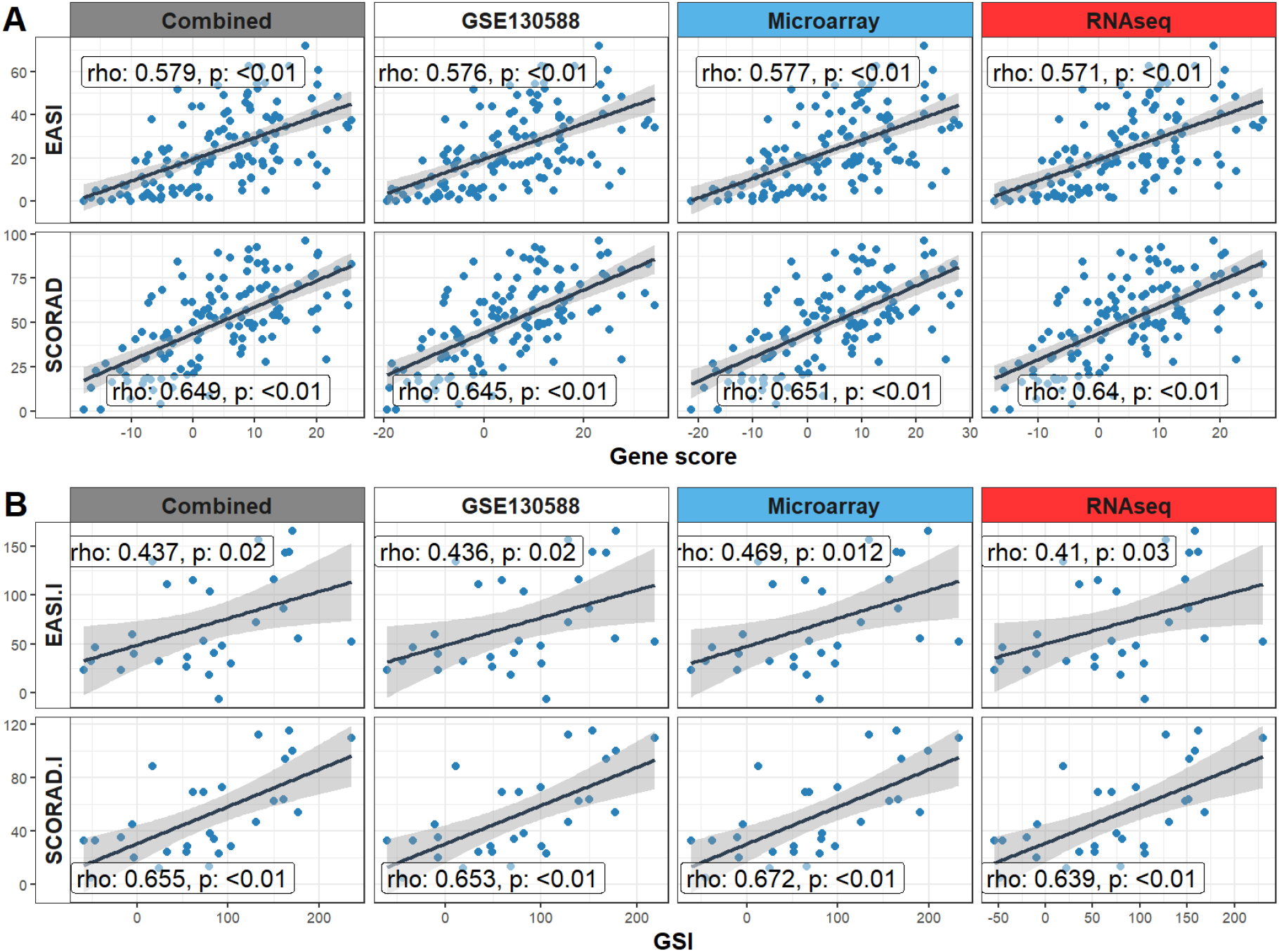
Disease severity index EASI and SCORAD correlation to gene scores from each meta-analyzed result and gene set obtained with one dataset (GSE130588) under the same threshold (|FC| *≥* 2, FDR *≤* 0.05) from published dataset (GSE130588).

**Supp. Fig S8.**
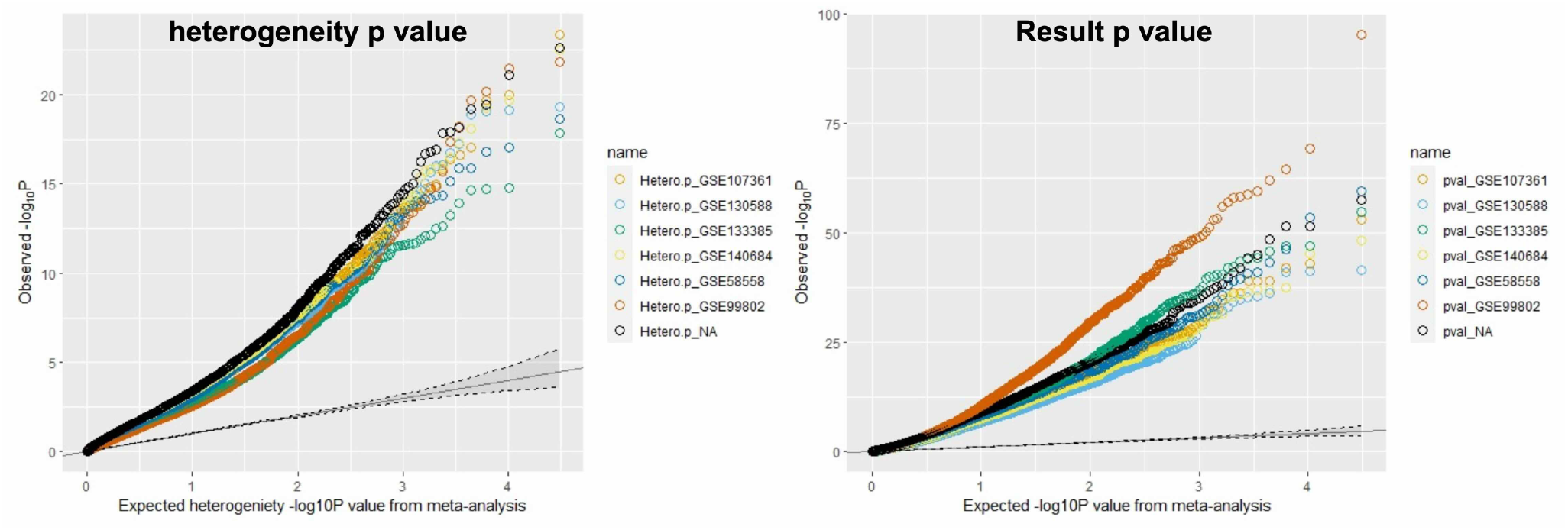
The Q-Q plot of leave-one-out analysis to remove one dataset at a time and process M-A with a fixed model to show the heterogeneity p-value and the result p-value. Each dotted curve shows the M-A result without that dataset, whereas on the right panel we can find the M-A result have a stronger statistical significance once GSE99802 has been removed.

**Table S1.** All datasets considered before applying inclusion criteria.

| Microarray datasets | RNA-seq datasets |
| --- | --- |
| E-MTAB-729 |  |
| GSE5667 |  |
| GSE60709 |  |
| GSE107361 |  |
| GSE32924 |  |
| GSE36842 | GSE121212 |
| GSE120899 | GSE137430 |
| GSE130588 | GSE140227 |
| GSE153007 | GSE157194 |
| GSE133385 | GSE141571 |
| GSE133477 |  |
| GSE140684 |  |
| GSE59294 |  |
| GSE27887 |  |
| GSE58558 |  |
| GSE99802 |  |

## Footnotes

1 https://www.metafor-project.org/doku.php/analyseskonstantopoulos2011#references

## Notes

### Competing Interest Statement

Drs. Li, He, Zhang, Page, Hyde, and Maciejewski are full-time employees of Pfizer and own stock or stock options in Pfizer.

### Summary of Updates

reformatted + moved supplementary doc to methods in the body of main ms

